# Matrix-matched calibration curves for assessing analytical figures of merit in quantitative proteomics

**DOI:** 10.1101/719179

**Authors:** Lindsay K Pino, Han-Yin Yang, William Stafford Noble, Brian C Searle, Andrew N Hoofnagle, Michael J MacCoss

## Abstract

Mass spectrometry is a powerful tool for quantifying protein abundance in complex samples. Advances in sample preparation and the development of data independent acquisition (DIA) mass spectrometry approaches have increased the number of peptides and proteins measured per sample. Here we present a series of experiments demonstrating how to assess whether a peptide measurement is quantitative by mass spectrometry. Our results demonstrate that increasing the number of detected peptides in a proteomics experiment does not necessarily result in increased numbers of peptides that can be measured quantitatively.

---

Mass spectrometry based proteomics has made great progress and is being used to address essential questions in basic biology and of biomedical significance. Of particular interest, the development of data independent acquisition mass spectrometry (DIA-MS) has made it possible to measure tens of thousands of peptides in a protein digest in 1-2 hours of instrument time. The sampling of tandem mass spectra in DIA-MS is unbiased [1] and systematic [2], in principle making it an appealing compromise between a narrowly focused targeted data acquisition strategy [3] and an irregularly sampled discovery method. Although fully targeted proteomics assays often include validation experiments to assess whether the change in measured signal is reflective of the actual change in peptide abundance, proteomics assays measuring thousands of analytes in an unbiased fashion rarely assess which peptide measurements are truly quantitative.

A measurement is quantitative when the change in measured signal reflects a change in the quantity of the analyte [4]. Specifically in mass spectrometry proteomics, for a method to be considered quantitative the relationship between the measured signal and the peptide quantity must be assessed. This assessment uses a *calibration curve*, where the analyte is diluted systematically to demonstrate that the measured signal is precise and above the *lower limit of quantitation* (LLOQ), the quantity below which a change in signal no longer reflects a change in quantity. Because liquid chromatography-tandem mass spectrometry is subject to matrix effects, calibration curves must be constructed in a relevant sample matrix. For endogenous compounds like peptides that are present in the sample matrix, assessment is frequently performed with *reverse calibration curves*, where a heavy isotope-labeled synthetic version of the analyte is diluted in the sample matrix [5, 6]. Although a signal measured below the LLOQ may still be used to assess a difference between two conditions, when compared to a signal above the LLOQ, the magnitude of the difference in signal is not reflective of the true difference in analyte quantity. In some papers, this phenomenon has been referred to as ratio compression [7]. Thus, unless the relationship between the quantity and signal for each analyte is documented, mass spectrometry measurements should be considered only differential rather than quantitative. In targeted proteomics studies, reverse calibration curves of increasing concentrations of stable isotope-labeled internal standard peptides can be used to approximate the LLOQ and precision of unlabeled peptide responses. However large-scale studies on the order of 1,000’s to 10,000’s of peptides like most DIA/SWATH-MS experiments do not evaluate peptide response. Calibration curves for up to 30 stable isotope-labeled internal standard peptides have been collected using DIA/SWATH-MS methods [8], but it is cost-prohibitive to synthesize stable isotope-labeled peptides for the number of targets detected in DIA. In this work, we propose a framework for discriminating between peptides that are only detectable and those which are both detectable and quantitative in a mass spectrometry experiment. We introduce an alternative to reverse calibration curves called *matrix-matched calibration curves*.

Our goal was to construct calibration curves and determine the LLOQ for every detectable peptide in a given complex protein mixture of interest using one dilution series and without pre-determining targets. We propose *matrix-matched calibration curves*, in which a complex protein sample of interest (a *reference material* [9]) is diluted with a *matrix-matched material*. A matrix-matched material may be any sample of equivalent biochemical complexity, but should not share any endogenous analytes with the reference material. For example, a matrix-matched material could be a stable-isotope labeled a reference material that preserve the matrix complexity but shift the peptide masses or using an equivalent biosample from an evolutionarily-diverged species (Supplemental Fig 1, 2). Each point in the dilution series has the same total protein concentration, composed of some ratio of the reference and matrix-matched material (Fig 1a) spanning several orders of magnitude (see Supplementary Table 1). A strength of this approach is that every peptide (or other type of analyte) in the reference material is diluted through the curve, meaning that calibration curves are constructed for all peptides detected in the reference material. To fit calibration curves to this novel data, we developed a computational model (Fig 1b) which extends the work described previously by Galitizine *et al*. [8] to accommodate the sparseness of matrix-matched calibration curve data and to determine the LLOQ for each detected analyte. Briefly, the model first fits a piece-wise linear regression to the noise and the signal segments of the curve data, then bootstraps the observed data, refits the piece-wise regression to the bootstrapped data to predict signal over the range of quantities measured. Finally we calculate the coefficient of variance (CV) of the predicted signal and define the LLOQ as the minimum quantity at which the predicted signal passes a predetermined CV threshold (CV ≤ 20% for the results reported here) (see Supplemental Methods).

**Figure 1:**
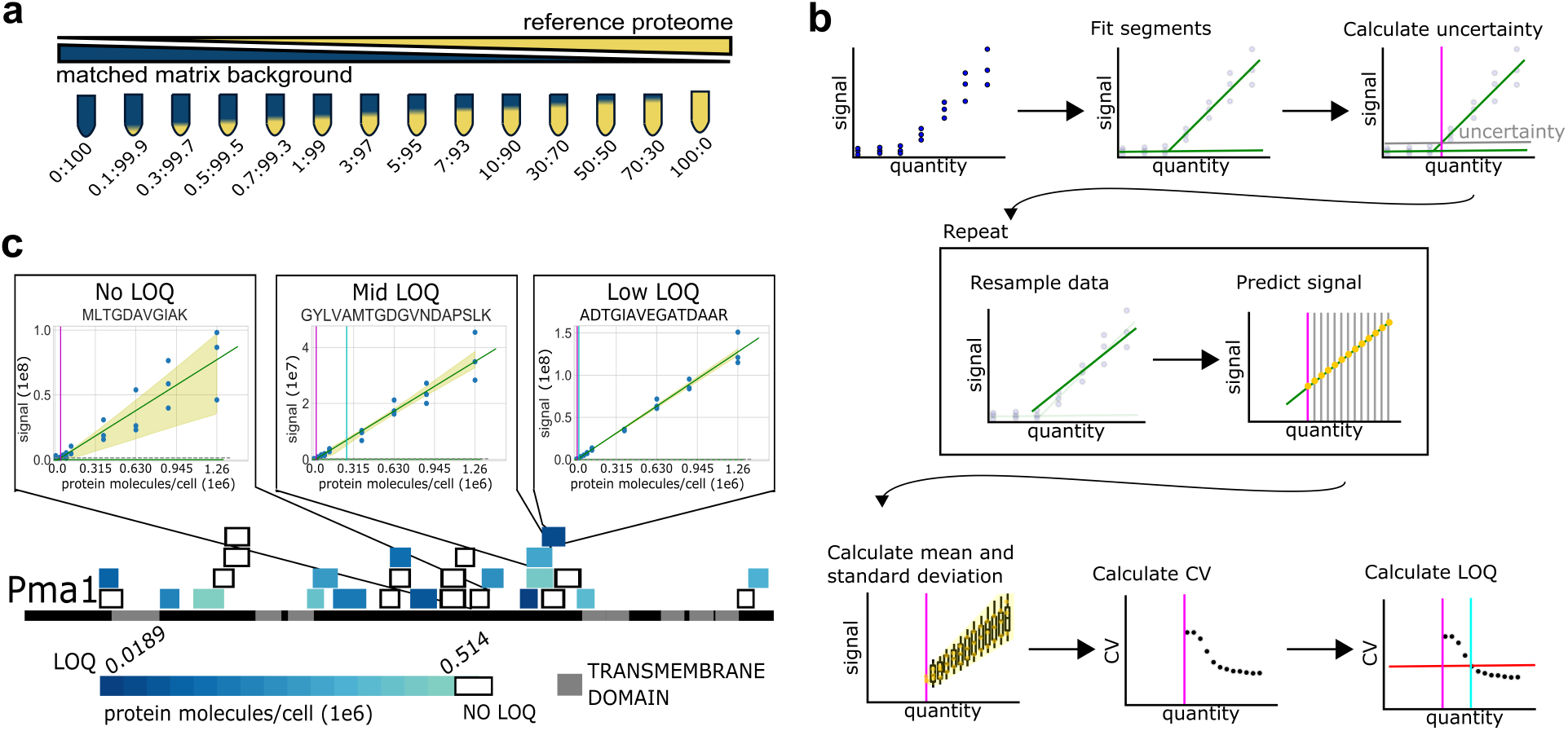
Constructing reference material calibration curves using a matched-matrix diluent. (a) A reference material is diluted into a matrix-matched material of similar matrix complexity but with no shared endogenous analytes, for example by stable isotope labeling the matrix or using a diverged species. The curve is made from dilutions spanning several orders of magnitude plus a *blank* with only the matrix-matched proteome. (b) The model for assessing the lower limit of quantification (LLOQ) using the sparse matrix-matched calibration curve data. We assess the LLOQ (cyan line) as the first point that is statistically different from the background (pink line) and has a CV *≤* 20% using bootstrapping (red line). (c) The sequence of plasma membrane ATPase (Pma1) is represented as a black line. The transmembrane domains along the sequence are depicted in grey. Each peptide detected by DIA-MS is represented by a colored box placed along the sequence. The color of the box ranks the peptide LLOQs. Three of the peptide calibration curves are shown above the sequence. Yellow shading indicates two standard deviations above and below the median for the bootstrapped data.

We apply the matrix-matched calibration curve framework first in yeast, and find that it high-lights the divide between detection and quantification especially at low protein abundances. In particular, highly abundant proteins often contain peptides that are detected at 1% FDR but are not quantifiable because the observed abundance in the reference material is below the LLOQ. Using the highly-abundant yeast proteome plasma membrane ATPase protein (Pma1) as an example, we detect 28 peptides at a 1% FDR threshold across the protein sequence (Fig 1c, Supplemental Fig 5-31). Of the detected peptides, only half (15 peptides) are deemed quantitative, and the quantitative peptides display a range of LLOQs spanning more than 20x. A peptide with no LLOQ is a less accurate quantitative proxy for Pma1, while a peptide with a low LLOQ is a more accurate quantitative proxy for Pma1 and is more accurate over a wider linear range. The extreme range of peptide responsiveness emphasizes the necessity to carefully select which peptides should act as quantitative proxies for their protein of origin.

The yeast proteome has the advantage of an established reference quantity for each protein, allowing us to contextualize our results. Ghaemmaghami *et al*. affinity-tagged the protein coding regions in yeast and reported the protein abundances in molecules-per-cell for 4,102 proteins, 3,869 of which could be quantified above 50 molecules/cell [10]. Using data independent acquisition mass spectrometry (DIA-MS) [11], we detected peptides from 2,870 of the proteins they quantified in the reference yeast proteome (Fig 2a, b). Using matrix-matched calibration curves to assess the quantitative accuracy of the detected peptides, we found that half of the detected proteins had at least one quantitative peptide (1,511 proteins). The proteins with validated peptides are primarily high quantity proteins, particularly those above 10,000 molecules/cell. As the reported quantity [10] decreases, fewer detected proteins have at least one quantitative peptide (Fig 2a, b). We compared the peptides determined to be quantitative by matrix-matched calibration curves with the peptides determined to be quantitative by a more conventional synthetic peptide approach [12]. Overall, the proposed framework assessed 6x more candidate peptides and defined 4.7x more peptides as quantitative (Supplemental Fig 3), demonstrating the higher throughput of the proposed framework compared to conventional approaches.

**Figure 2:**
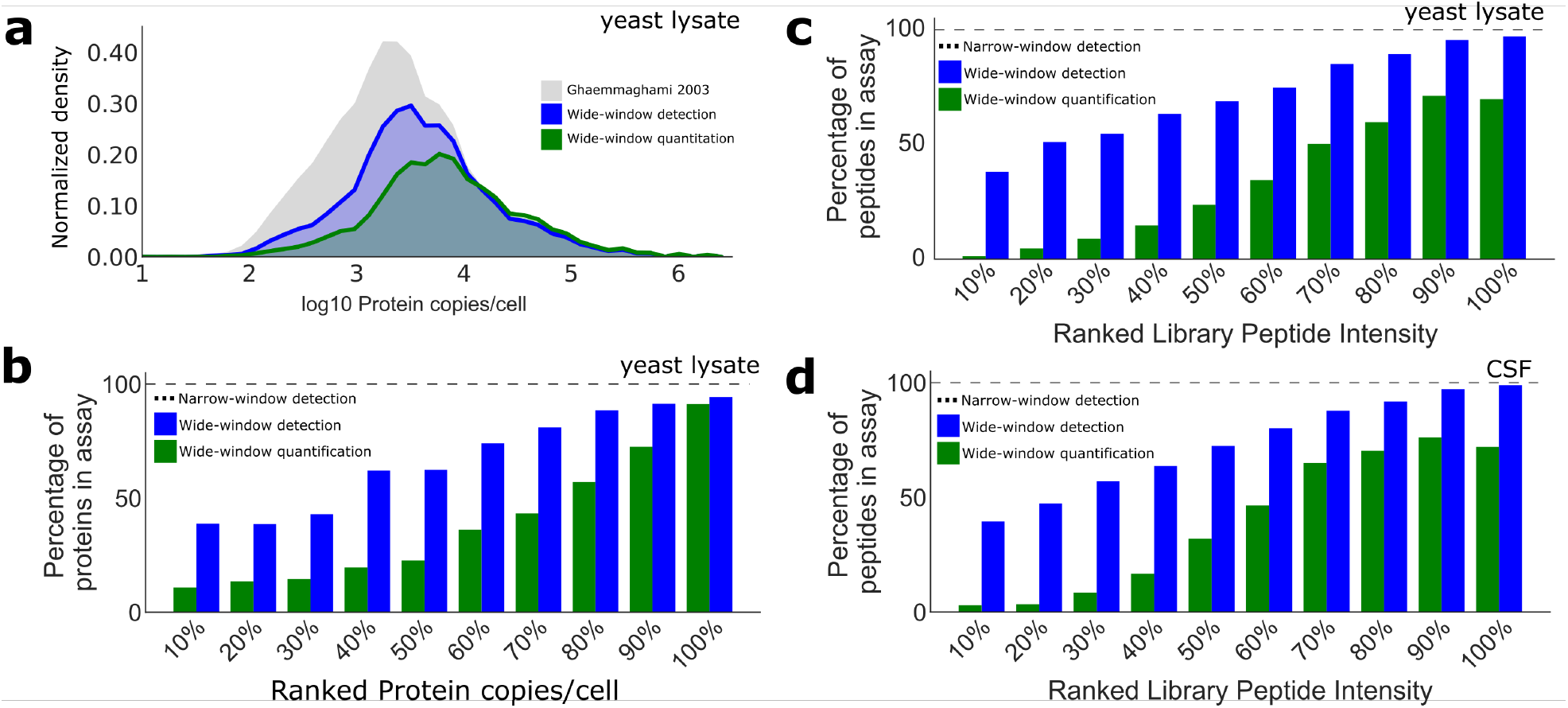
There is a difference between the detection of a peptide and the quantification of a peptide. The (a) number and (b) percentage of proteins detected in yeast at different orders of magnitude of abundance. Ghaemmaghami *et al*. comprehensively estimated protein copies per cell in yeast (black, 3,869 proteins) using epitope tagging [10]. The wide-window DIA using a chromatogram-library approach [11] detects, at 1% protein-level FDR, 74% of these proteins (blue, 2,870 proteins). The number of proteins quantifiable by DIA-MS (proteins with at least one peptide with a defined LLOQ) encompasses 52% of the detected proteins, or 39% of the expressed proteins (green, 1,511 proteins). (c) Peptides detected in the yeast lysate narrow-window library are ranked by intensity, and the wide-window detected and quantitative peptides are shown for each decile. (d) Cerebrospinal fluid peptides detected in the narrow-window library (8,698 total peptides, 2,994 protein groups) are ranked by intensity, and the wide-window detected and quantitative (3,183 peptides; 1,303 protein groups) peptides are shown for each decile.

The matrix-matched calibration curve approach is generalizable beyond cell culture. To illustrate its flexibility, we adapted the framework to two human samples: cerebrospinal fluid (CSF) and formalin-fixed paraffin embedded (FFPE) tissue. For the CSF reference material, we chose a commercially-available pool of healthy donor CSF (Golden West Biologicals, Inc.) which we prepared following conventional protocols. For the CSF matrix-matched material, we performed a second enzymatic digest in the presence of ^18^O-enriched water. This reaction preferentially exchanges one or both of the oxygens at the C-terminus of the peptide with ^18^O, shifting the peptides by 2 or 4 mass units via incorporation of one or two ^18^O atoms. Following the matrix-matched calibration curve framework, we found that 36% of peptides detected in the CSF reference material library (8,698 peptides; 2,994 protein groups) have a defined LLOQ (3,183 peptides; 1,303 protein groups) (Fig 2d). In both the yeast (Fig 2c) and CSF (Fig 2d) references, the most intense peptides in the reference are more likely to be detected and quantified. We also applied the matrix-matched calibration curve approach to an FFPE sample (Supplemental Fig 4) and acquired the data by another form of mass spectrometry (selected reaction monitoring). To construct the FFPE matrix-matched calibration curve, we spiked human plasma into homogenized chicken liver as a reference and used the unspiked homogenized chicken liver for the background proteome. We targeted 84 peptides (18 proteins) and found that 27 of the targeted peptides were quantitative (14 proteins). This demonstrates that the matched-matrix calibration curve approach is generalizable broadly across not only sample types but also mass spectrometry acquisition approaches.

A limitation of the approach is that the maximum possible peptide quantification is limited by the endogenous abundance of the peptide in the reference, which for low abundance peptides results in stunted linear range. Another consequence of the endogenous abundance limitation is that matrix-matched calibration curve data is extremely sparse compared to conventional calibration curves because low abundance reference peptides produce low signal which reduces to zero signal as the reference is diluted. Additionally, while the quantitative peptides reported here may serve as a starting point for future assay development, we emphasize that these LLOQs are specific to these exact conditions. Matrix-matched calibration curves, like all calibration curves, are only reflective of the peptide measured on a given platform. While most quantitative methods report precision, this does not assess whether a change in signal reflects the change in quantity. Therefore, the use of matrix-matched calibration curves should be performed for all proteomics experiments that require an assessment of which peptides reflect the change in quantity those that are just differential.

