## Supplement for "Matrix-matched calibration curves for assessing analytical figures of merit in quantitative proteomics"

### MATRIX-MATCHED CALIBRATION CURVES SUPPLEMENT

Lindsay K Pino
Department of Genome Sciences
University of Washington

Brian C Searle
Department of Genome Sciences
University of Washington

Han-Yin Yang
Department of Genome Sciences
University of Washington

Andrew N Hoofnagle
Department of Laboratory Medicine
University of Washington

William Stafford Noble
Department of Genome Sciences
Department of Computer Science and Engineering
University of Washington

Michael J MacCoss
Department of Genome Sciences
University of Washington

July 30, 2019

### 1 Supplemental Methods

#### 1.1 Sample preparation and mass spectrometry data acquisition.

**Yeast culture and sample preparation.** Yeast strains BY4741 (MATa his3Δ1 leu2Δ0 met15Δ0 ura3Δ0) and S288C (MATα) (Dharmacon) were cultured in YEPD and $^{15}$N minimal media, respectively, for matched matrix calibration curve experiments. Cultures of 50 mL were grown to mid-log phase, harvested, and lysed individually with 8M urea buffer solution and bead beating (7 cycles of 4 minutes beating with 1 min rest on ice). Cell lysates were reduced with 5mM DTT, alkylated with 15mM IAA, and digested for 16 hours with 1:50 trypsin to protein. The peptide digests were desalted with a mixed-mode (MCX) method, dried down via speedvac overnight, and brought up with synthetic iRT peptide standards (Pierce Peptide Retention Time Calibration Mixture) to 1$\mu$g/$\mu$l total proteome using calculations from a bicinchoninic acid (BCA) assay (Pierce BCA Protein Assay Kit) performed on the lysate.

**Cerebrospinal fluid sample preparation.** Pooled human cerebrospinal fluid (CSF) from healthy donors was purchased from Golden West Biologicals. CSF was denatured with 0.2% PPS Silent Surfactant, reduced with 5mM DTT, alkylated with 15mM IAA, and digested for 16 hours with 1:25 trypsin to protein. The peptide digest were desalted with a mixed-mode (MCX) method, the desalted peptides split into two aliquots, and each aliquot dried down via speedvac overnight. 24 hours prior to MS acquisition, one dried aliquot was resuspended in 0.05$\mu$g/$\mu$L trypsin in $^{18}$O-enriched water (purchased from Cambridge Isotope Laboratories, Inc.) following a standard $^{18}$O-labeling protocol [4], the other was resuspended in 0.05$\mu$g/$\mu$L trypsin in conventional molecular-grade water. The digest incubated overnight then was quenched with 5mM DTT, cooled to room temperature, and acidified with formic acid.

**Formalin-Fixed Paraffin-Embedded (FFPE) sample preparation.** Pooled human plasma (75$\mu$g/$\mu$l; Na-Citrate, Cat 7303806, Unit 23-45456A) were diluted with DPBS (Life technologies,

14190-144) to make a plasma dilution series with 13 different concentrations. The 30$\mu$l of human plasma or blank samples was well mixed with 80$\mu$l homogenized chicken liver in an open-ended syringe (company name and size). Each concentration mixture was quickly mixed with 200$\mu$l 20% formalin and followed by 90$\mu$l 1% agarose. The syringe was then sealed and left on the bench overnight at room temperature to allow protein-liver mixture form a gel-like structure. Each resulting product was then pushed out from syringe gently and placed into a tissue cassette for standard paraffin embedding procedure.

Six of 10$\mu$m-thick tissue slides were obtained from each protein-chicken liver block, and then deparaffined. Proteins on the deparaffinized tissue slides were re-solubilized in 60$\mu$l 0.1% RapiGest buffer by undergoing high heat and sonication cycles. Reconstituted protein mixture was reduced, alkylated and digested with 5 $\mu$l trypsin overnight. The protein digests were stored in -80 C until the day of analysis.

**Liquid chromatography mass spectrometry.** Peptides were separated by liquid chromatography before analysis by mass spectrometry, either with a Waters NanoAcquity UPLC for yeast and human CSF DIA experiments or a Thermo easy-nanoLC for FFPE tissue block SRM experiments. On all systems, peptides were separated by reverse phase liquid chromatography using pulled tip columns created from 75 $\mu$m inner diameter fused silica capillary (New Objectives, Woburn, MA) in-house using a laser pulling device and packed with 3 $\mu$m ReproSil-Pur C18 beads (Dr. Maisch GmbH, Ammerbuch, Germany) to 30 cm. Trap columns were created from 150 um inner diameter fused silica capillary fritted with Kasil on one end and packed with the same C18 beads to 3 cm.

*$^{14}$N BY4741 yeast proteome separation on Waters NanoAcquity UPLC.* Solvent A was 0.1% formic acid in water (v/v), solvent B was 0.1% formic acid in 98% acetonitrile (v/v). For each injection, approximately 1 $\mu$g total protein was loaded and eluted using a 90-minute gradient from 5 to 35% B, followed by a 40 minute wash and equilibration (35 to 60% B for 10 minutes, 60 to 95% B for 5 minute, 95% B for 5 minutes, 95 to 2% B for 1 minute, 2% B for 19 minute).

*$^{16}$O human CSF proteome separation on Waters NanoAcquity UPLC.* Solvent A was 0.1% formic acid in water (v/v), solvent B was 0.1% formic acid in 98% acetonitrile (v/v). For each injection, approximately 1 $\mu$g total protein was loaded and eluted using 60-minute gradient from 5 to 35% B, followed by a 40 minute wash and equilibration (35 to 60% B for 10 minutes, 60 to 95% B for 5 minute, 95% B for 5 minutes, 95 to 2% B for 1 minute, 2% B for 19 minute).

*FPE tissue block proteome separation on Thermo easy-nanoLC.* Solvent A was 0.1% formic acid in water (v/v), solvent B was 0.1% formic acid in 98% acetonitrile (v/v). For each injection, approximately 1 $\mu$g total protein was loaded and eluted using a 30-minute gradient from 0 to 40% B, followed by a 18 minute wash and equilibration (40 to 60% B for 5 minutes, 60% B for 5 minutes, 60 to 100% B for 1 minute, 100% B for 5 minutes, 100 to 0% B for 1 minute, 0% B for 1 minute).

**Data independent acquisition mass spectrometry (DIA-MS).** Yeast curve data were acquired using data-independent acquisition (DIA) method on a Thermo Q-Exactive HF Orbitrap mass spectrometer. Human CSF curve data were acquired using an equivalent DIA method on a Thermo Lumos mass spectrometer. Both DIA methods followed the chromatogram library workflow, described in greater detail elsewhere [5]. Briefly, to create the chromatogram library, the mass spectrometer was configured to acquire six gas phase fractions of the undiluted reference proteome for each curve (e.g. $^{14}$N BY4741 yeast proteome, $^{16}$O human CSF).

*Thermo Q-Exactive HF Orbitrap method details.* Mass range of 388.43190-1,012.70480 m/z was monitored in the yeast experiments. The chromatogram library, gas-phase fractionated "narrow window" Thermo QEHF method details were as follows: 4 m/z overlapped windows (effectively 2 m/z isolation), 30k resolution, 55 maximum ion inject time, 1e6 AGC. The quantitative, single-shot "wide window" Thermo QEHF method details were as follows: 24 m/z overlapped windows (effectively 6 m/z isolation), 30k resolution, 55 maximum ion inject time, 1e6 AGC. All DIA spectra were programmed with a normalized collision energy of 27 and an assumed charge state of +2.

*Thermo Lumos method details.* Mass range of 394.4319-1,006.704807 m/z was monitored in the CSF experiments. The chromatogram library, gas-phase fractionated "narrow window" Thermo Lumos method details were as follows: 4 m/z overlapped windows (effectively 2 m/z isolation), 30k resolution, 60 maximum ion inject time, 4e5 AGC. The quantitative, single-shot "wide window" Thermo Lumos method details were as follows: 12 m/z overlapped windows (effectively 6 m/z isolation), 15k resolution, 20 maximum ion inject time, 4e5 AGC. All DIA spectra were programmed with a normalized collision energy of 33%.

Thermo RAW files were converted to .mzML format using the ProteoWizard package (version 3.0.10106), where they were peak picked using vendor libraries. Converted acquisition files were processed using EncyclopeDIA (version 0.8.0) configured with default settings (10 ppm precursor, fragment, and library tolerances, considering both B and Y ions, and trypsin digestion was assumed). EncyclopeDIA was configured to use Percolator (version 3.1).

**Selected reaction monitoring mass spectrometry (SRM-MS).** FFPE tissue block curve data were acquired using a targeted SRM-MS method on a Thermo TSQ Quantiva triple quadrupole mass spectrometer. The target list was developed based on clinical relevance to amyloidosis. Instrument details were as follows: dwell time 2ms, Q1 resolution set to 0.7 FWHM, Q3 resolution set to 0.7 FWHM, CID gas set to 1.5 mTorr.

**Data availability.** The RAW files, converted MZML files, Encyclopedia elib files, and Skyline documents have been deposited in ProteomeXChange Consortium [7] via the Panorama [6] partner repository with the identifiers PXD014815 (ProteomeXchange) and `https://panoramaweb.org/matrix-matched_calcurves.url` (Panorama).

#### 1.2 Constructing a serial dilution standard curve using the matched matrix calibration curve approach.

The curves used in this work followed Clinical and Laboratory Standards Institute (CLSI) recommendations [3]. Specifically, the CLSI recommends calibration curves for LC-MS assays are composed of at minimum a blank (a sample containing matrix only) and six to eight calibration standards, with the calibration standards commonly spaced logarithmically across several orders of magnitude. Our yeast dilution series is composed of 13 calibration points and a blank consisting of the matched matrix alone (Supplemental Table 1). It is also recommended that calibration curves not be composed of one continuous serial dilution, because this can propagate pipetting errors throughout the curve. We therefore constructed these yeast calibration curves as a set of five serial dilutions, with each of points A, B, C, D, and E mixed individually from reference and matched matrix materials, then subsequent points are dilutions of those original five (F is a dilution of B, G is a dilution of C, H is a dilution of D, I is a dilution of E; then J is a dilution of F, K is a dilution of G, L is a dilution of H, and M is a dilution of I). If pipetting error occurred in one of the dilutions, it would appear as an outlying point in the final calibration curve.

For the cerebrospinal fluid curves, we followed the same fractional dilution scheme as above, but did not include points K, L, and M due to limited availability of the matched matrix material ($^{18}O$ enriched CSF).

| Point | Yeast reference (fractional dilution) | Yeast matched matrix (fractional dilution) |
|---|---|---|
| A | 1 | 0 |
| B | 0.7 | 0.3 |
| C | 0.5 | 0.5 |
| D | 0.3 | 0.7 |
| E | 0.1 | 0.9 |
| F | 0.07 | 0.93 |
| G | 0.05 | 0.95 |
| H | 0.03 | 0.97 |
| I | 0.01 | 0.99 |
| J | 0.007 | 0.993 |
| K | 0.005 | 0.995 |
| L | 0.003 | 0.997 |
| M | 0.001 | 0.999 |
| N | 0 | 1 |

Table 1: Dilution series for the yeast matched matrix calibration curves. Using a fractional dilution scheme, the reference material is diluted with the matched matrix to create a dilution series. In this 14-point design, the calibration standards span three orders of magnitude of the reference material and include a blank with only the matched matrix.

| Point | Plasma (fractional dilution) | PBS (fractional dilution) |
|---|---|---|
| A | 1 | 0 |
| B | 0.5 | 0.5 |
| C | 0.25 | 0.75 |
| E | 0.1 | 0.9 |
| F | 0.05 | 0.95 |
| G | 0.025 | 0.975 |
| I | 0.01 | 0.99 |
| J | 0.005 | 0.995 |
| K | 0.0025 | 0.9975 |
| L | 0.00125 | 0.99875 |
| M | 0.001 | 0.999 |
| O | 0 | 1 |

Table 2: Dilution series for the FFPE tissue block matched matrix calibration curves. Using a fractional dilution scheme, healthy donor plasma (reference) is diluted with PBS to create a dilution series. An equal volume of each calibration point was then mixed with a homogenate of chicken liver and prepared as individual FFPE tissue blocks.

For the FFPE tissue block proof of concept, we created concentration points of human plasma by diluting healthy donor pooled plasma into PBS, then mixing an equal volume of each plasma dilution with liver homogenate using an open-end 2ml syringe (Supplemental Table 2). Each concentration point-spiked liver homogenate sample was then formalin fixed and paraffin embedded into individual tissue blocks. Tissue blocks were scraped and prepared for analysis by mass spectrometry as described above.

##### 1.3 A piecewise linear model to fit sparse, label-free LC-MS calibration curves.

We developed a model to fit the data produced by the matched matrix calibration curve method. The model is an extension of the work described previously by Galitizine et al [1]. Below, we briefly summarize the main steps of the model then discuss each step in detail.

**Algorithm 1** Model for determining LOD and LOQ from matched matrix calibration curves

```
Input: x curve points, y measured signals.
Fit piecewise regression (parameters b_n, b_s, m_s)
Find intersection of piecewise components (P_x = (b_n − b_s)/m_s)
Calculate standard deviation of noise segment (σ_{y_n})
Calculate LOD (LOD = (b_n + σ_{y_n} − b_s)/m_s)
Uniformly discretize 100 bins of x_i from the range LOD < x_max
for i to N do
    Resample n = x data points from x, y with replacement
    Fit piecewise regression to the resampled points
    For each x_i predict y_i using the regression parameters
    For each x_i calculate CV_{y_i} = σ_{y_i}/μ_{y_i}
end for
Calculate LOQ (LOQ = min(x_i) for which CV_{y_i} ≤ 0.2)
```

First, the model assumes two segments are present in the calibration curve: a noise segment where the measured signal $y_n$ (reported as intensity, peak area, estimated concentration, etc) does not exceed background noise and a signal segment where the measured signal $y_s$ is within the linear range for the analyte. Formally, we express this model in Equation 1 as

$$f(x) = \begin{cases} y_n = b_n & x < LOD \\ y_s = m_s x + b_s & x > LOD \end{cases} \tag{1}$$

where $x$ is the experimentally constructed analyte dilution values given by concentration, copies-per-cell, fractional dilution, etc. We use weighted least squares to minimize the function (lmfit package version x.x) using as weights the inverse square root of the curve points, and we constrain the parameters $(b_n, b_s, m_s)$ following (Equation 2)

$$\begin{aligned} m_s &\geq 0 \\ b_n &\geq b_s \\ b_n &\geq 0 \end{aligned} \tag{2}$$

With these constraints, we enforce that the signal segment must have a positive, nonzero slope, and we enforce that the intersection of the noise and the signal segments must be positive.

To determine the standard deviation associated with the noise segment, we calculate the empirical standard deviation of all $y_n$ values associated with the noise segment. The $y_n$ values are those where the corresponding $x_n$ values are less than the intersection $P_x$ of the noise and linear segments.

$$P_x = \frac{b_n - b_s}{m_s} \tag{3}$$

Thus, we compute the empirical standard deviation $\sigma_{y_n}$ in $y_n$ for all points for which $x \leq P_x$.

Next we determine the figures of merit: limit of detection (LOD) and limit of quantitation (LOQ). We define the limit of detection (LOD) as the $x$ for which the corresponding signal $y$ is one standard deviation ($\sigma$) above the noise segment,

$$LOD = \frac{b_n + \sigma_{y_n} - b_s}{m_s} \tag{4}$$

The limit of quantitation (LOQ; also referred to as the Lower Limit of the Measuring Interval (LLMI)) is defined by the Clinical and Laboratory Standards Institute [3] as ”the lowest measurand concentration at which all defined performance characteristics of the measurement procedure are met.” The performance characteristics we choose to define are the lowest analyte concentration which (1) is above the LOD and (2) achieves a coefficient of variation (CV) less than a threshold $\tau$ selected by a researcher (default is a 20% CV, $\tau = 0.2$). To determine the value $x$ which meets these two criteria, we first uniformly discretize the range of $x$ above the LOD into 100 bins ($x_i$), for which we will calculate 100 predicted $y_i$ by bootstrapping. Then we calculate the standard deviation and mean in the 100 predicted $y_i$ for each $x_i$.

For bootstrapping, we resample the entire dataset with replacement $N$ times (default $N = 100$). Each of the $N$ resampled data sets is fit to the piecewise regression model described in 1. We use the piecewise regression parameters to calculate the predicted response for a series of curve points spanning the range of curve points in the empirical data. The mean and standard deviation of the bootstrapped $y_i$ values are used to calculate a bootstrapped coefficient of variance ($CV_{y_i}$) for each of the curve points in the series. Last, the LOQ is calculated as the lowest value in the curve point series above the LOD which passes at or below the $CV_{y_i}$ threshold specified by the researcher (default $CV_{y_i} = 0.2$). The user has the option to set more or less conservative thresholds. For instance, the CV threshold recommended by Clinical and Laboratory Standards Institute guidelines is 10:1 signal:noise which equates to a $CV_{y_i} = 0.1$ threshold [3].

The code is accessible on Bitbucket (`https://bitbucket.org/lkpino/matrix-matched_calcurves`).

#### 2 Supplemental Figures

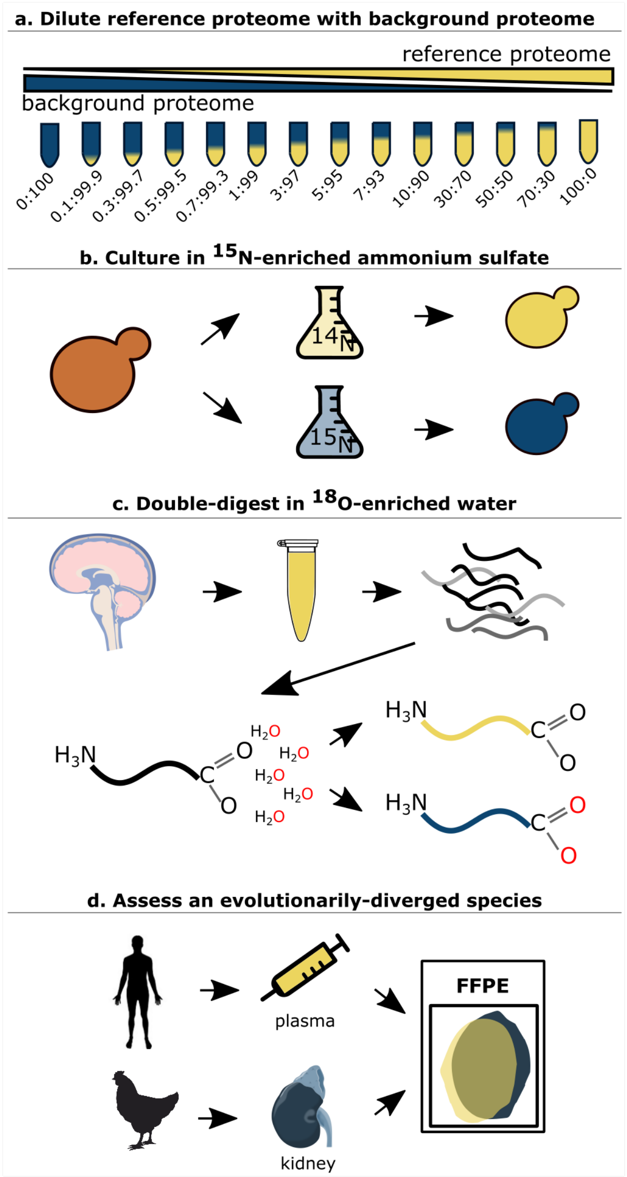


Figure 1: Three methods for constructing matched matrix calibration curves from a reference proteome. (a) All matched matrix calibration curves are constructed from a reference proteome and a background proteome. The reference proteome is diluted into a background proteome spanning several orders of magnitude in ratio. (b) Cell lysates can be matrix-matched by culturing the cells in a stable isotope media, such as $^{15}$N or SILAC, in order to shift m/z of the background matrix. (c) Biofluids such as cerebrospinal fluid or plasma can me matrix-matched by incorporating $^{18}$O into peptides via trypsin incubation in $^{18}$O-enriched water. (d) Tissues or other biological samples requiring sensitive preparation protocols can be matrix-matched using an evolutionarily-diverged species, ensuring that the matrix is similarly complex but contains no homologous analytes.

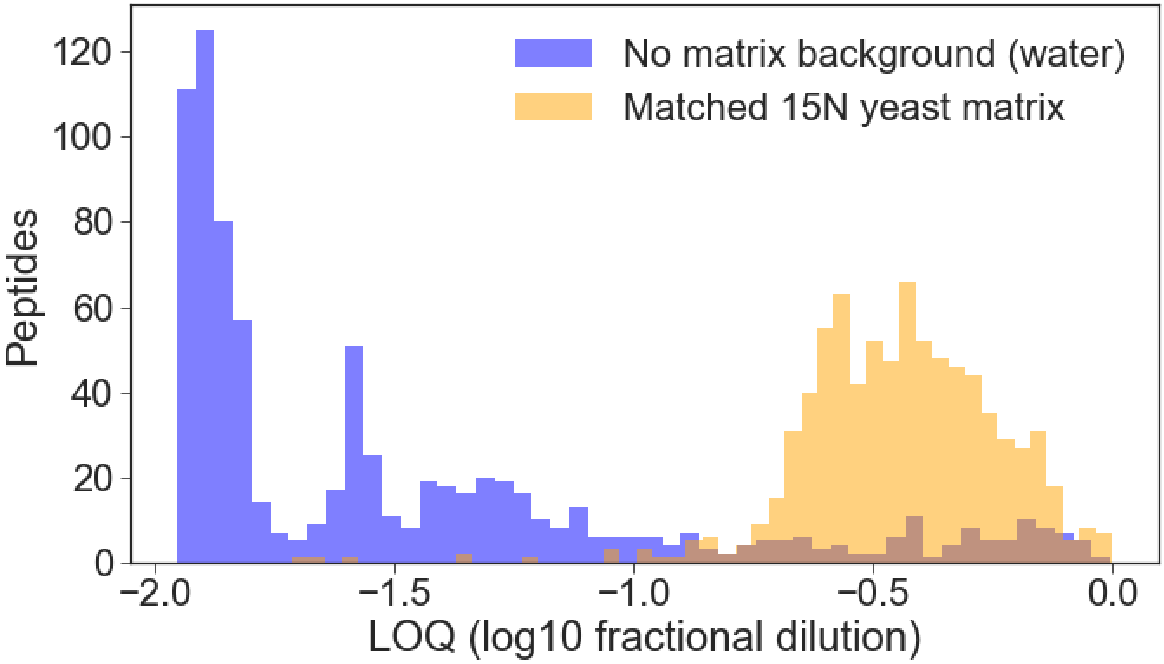


Figure 2: Reference materials must be diluted with a similarly complex material to preserve matrix properties. The yeast reference material was diluted in a matched matrix with 15N-shifted yeast (orange) or diluted in water with no matrix replacement (blue). Our curve-fitting model was then run on each data set to calculate the LOQs. The reported peptide LOQs are significantly more sensitive when the matrix complexity is not replaced, showing the importance of retaining matrix properties when building the calibration curve.

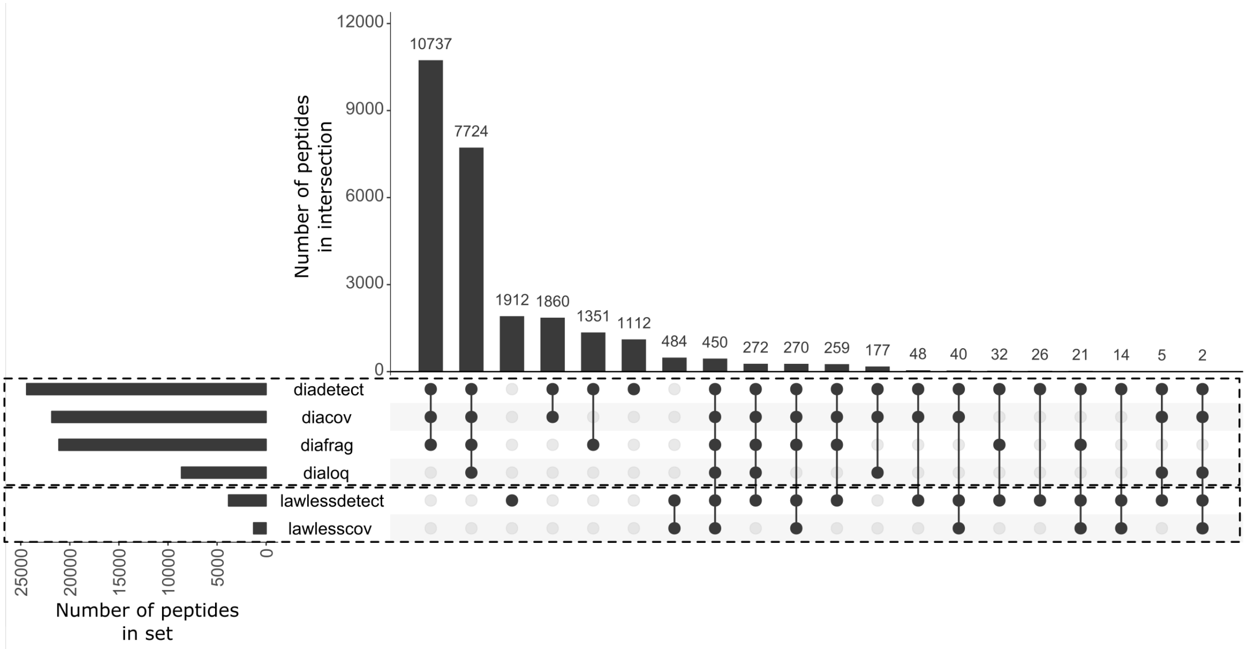


Figure 3: Matched matrix calibration curves can assess more candidate targets than conventional approaches without predetermining targets. The UpSet plot compares yeast peptides detected and deemed quantitative by DIA-MS and matched matrix calibration curves (this work, legend ”dia*”) with the peptides detected and validated by QConCat SRM-MS in Lawless et al 2016 [2]. Of all peptides detected in the wide-window DIA-MS at 1% FDR (”diadetect”, 24,400 peptides), only 6,117 peptides (25%) display all three desirable quantitative traits (”diacov”, peptides with ≤20% CV in the undiluted yeast sample; ”diafrag”, peptides with ≥3 interference-free fragment ions; ”dialoq”, peptides with a defined LOQ). Lawless et al 2016 assessed over 4,000 total peptides (”lawlessdetect”, QConCat peptides tested by Lawless et al 2016), of which 1,281 peptides (50%) displayed the desired quantitative properties (”lawlesscov”, QConCat peptides tested by Lawless 2016 with ≤ 20% CV). Overall, the proposed framework assessed 6x more candidate peptides and defined 4.7x more peptides as quantitative. The quantitative peptides in the proposed approach map to 1,629 proteins; the quantitative peptides by the QConCat approach map to 644 proteins. Both approaches include quantitative peptides for 520 proteins, and the QConCat approach includes peptides for 124 proteins not represented by the proposed approach, while the proposed approach includes 1,109 proteins not represented by the QConCat approach.

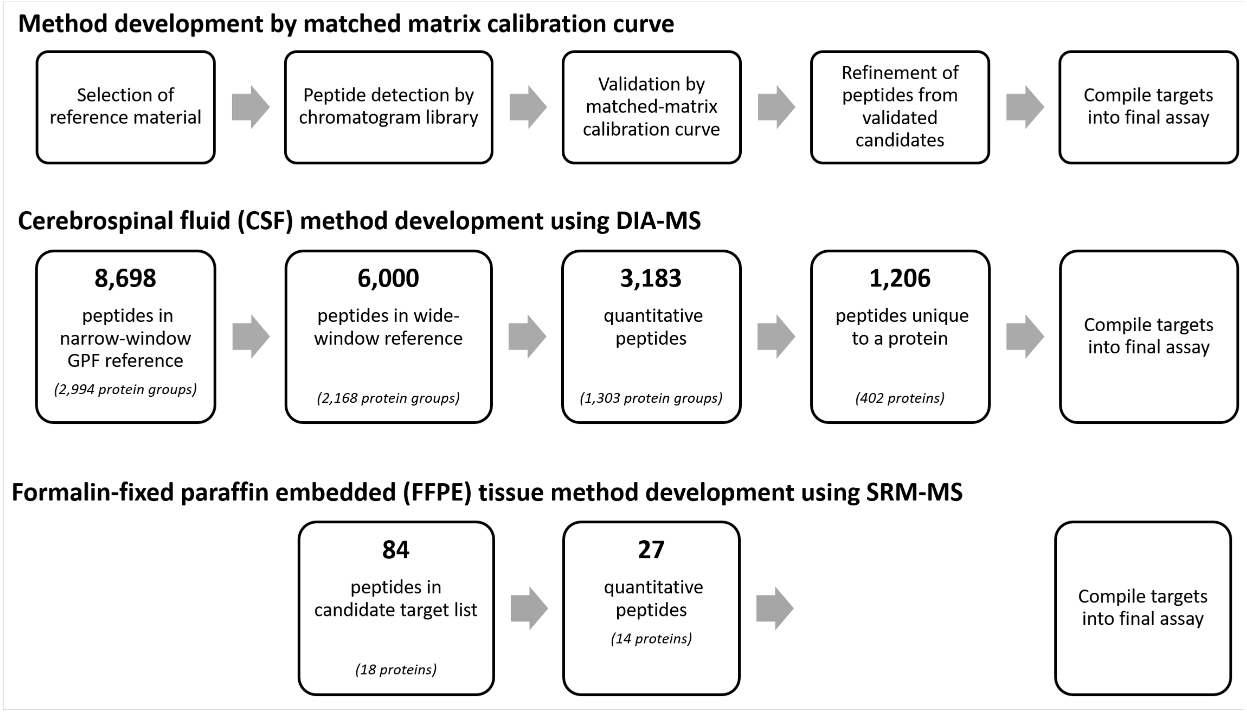


Figure 4: Matched matrix calibration curves can be used to rapidly develop targeted methods. Starting with all peptides detected in a gas-phase fractionated reference material as possible candidates, the first refinement step in this protocol discards any candidate target that cannot be detected in an unfractionationated, single-shot acquisition of the reference material. Next, the matched matrix calibration curve framework is used to assess quantitative figures of merit, discarding any candidates whose abundance in the reference material is below the analyte LOQ. For most targeted quantitative proteomics work, targets that are unique to a protein are considered better candidates. Even if experimenters are not starting from DIA data, the matched matrix calibration curve approach can be used to quickly eliminate poor quantitative targets from an assay.

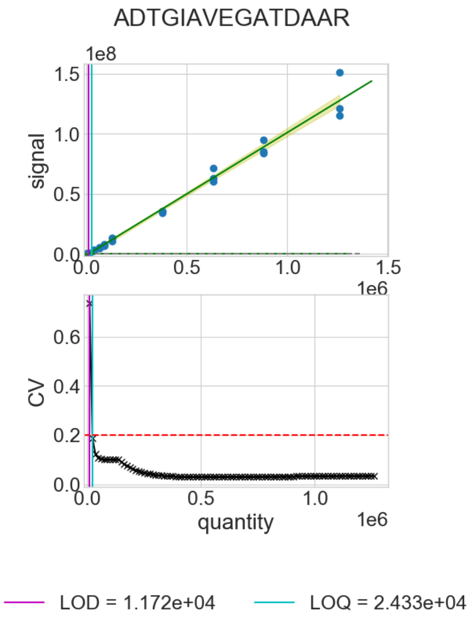


Figure 5: Calibration curve model outputs for Pma1 peptide ADTGIAVEGATDAAR.

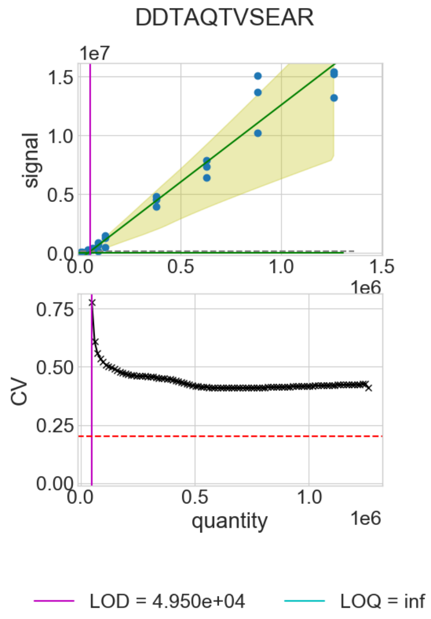


Figure 6: Calibration curve model outputs for Pma1 peptide DDTAQTVSEAR.

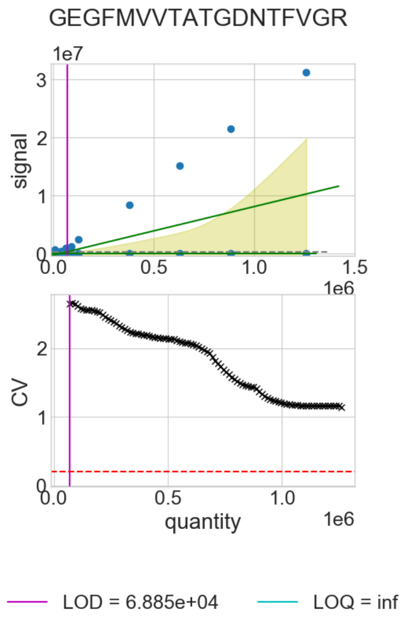


Figure 7: Calibration curve model outputs for Pma1 peptide GEGFMVVTATGDNTFVGR.

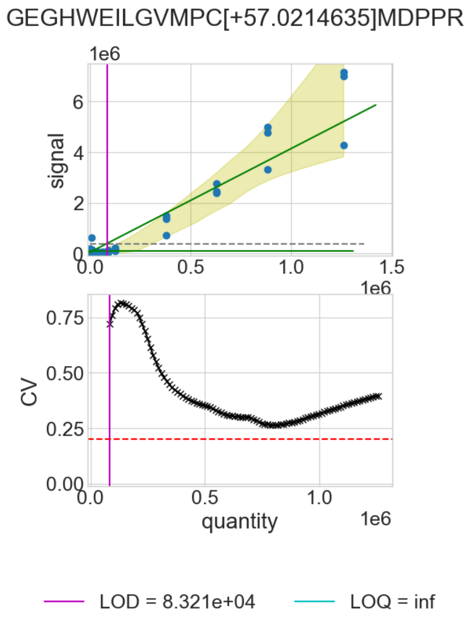


Figure 8: Calibration curve model outputs for Pma1 peptide GEGHWEIL-GVMPC[+57.0214635]MDPPR.

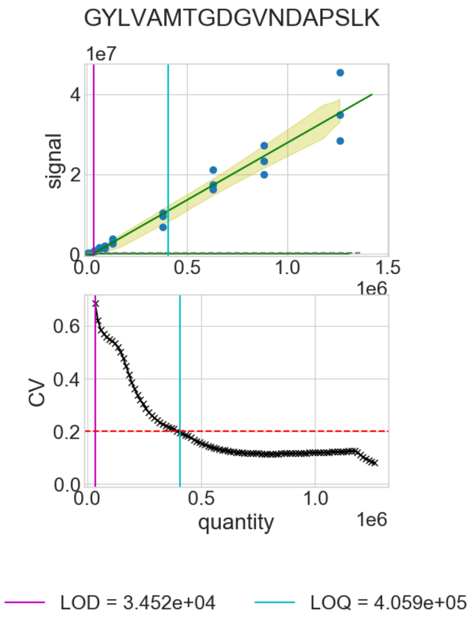


Figure 9: Calibration curve model outputs for Pma1 peptide GYLVAMTGDGVNDAPSLK.

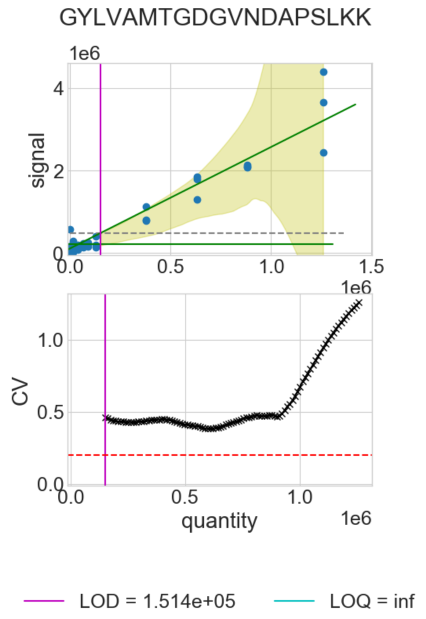


Figure 10: Calibration curve model outputs for Pma1 peptide GYLVAMTGDGVNDAPSLKK.

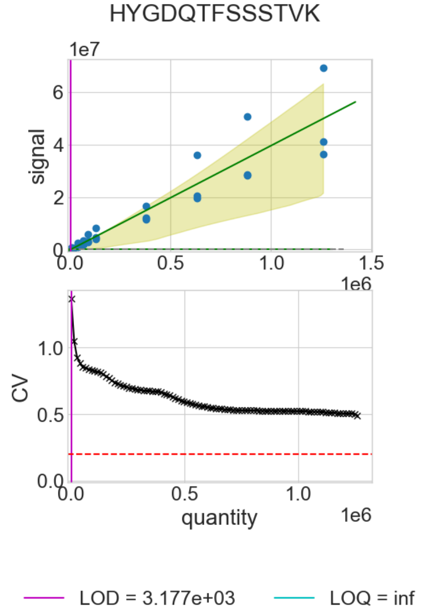


Figure 11: Calibration curve model outputs for Pma1 peptide HYGDQTFSSSTVK.

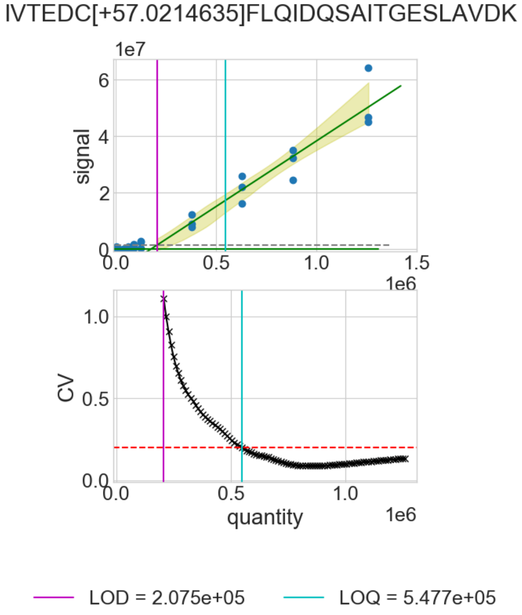


Figure 12: Calibration curve model outputs for Pma1 peptide IVTEDC[+57.0214635]FLQIDQSAITGESLAVDK.

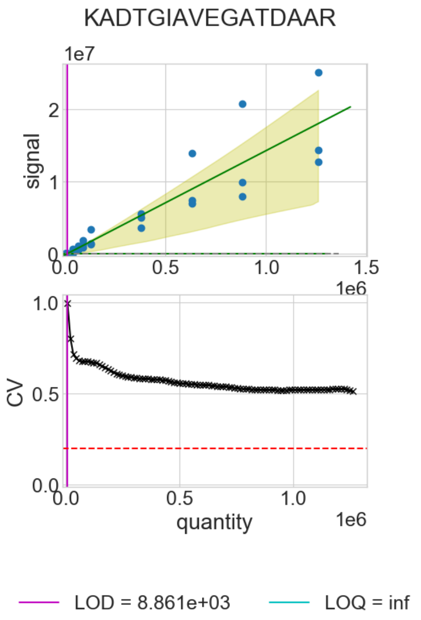


Figure 13: Calibration curve model outputs for Pma1 peptide KADTGIAVEGATDAAR.

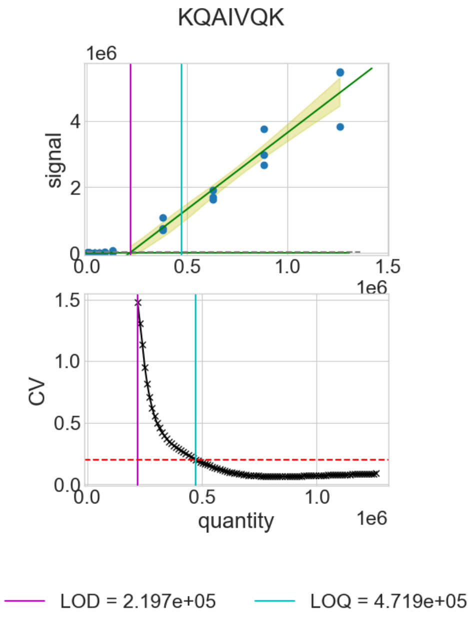


Figure 14: Calibration curve model outputs for Pma1 peptide KQAIVQK.

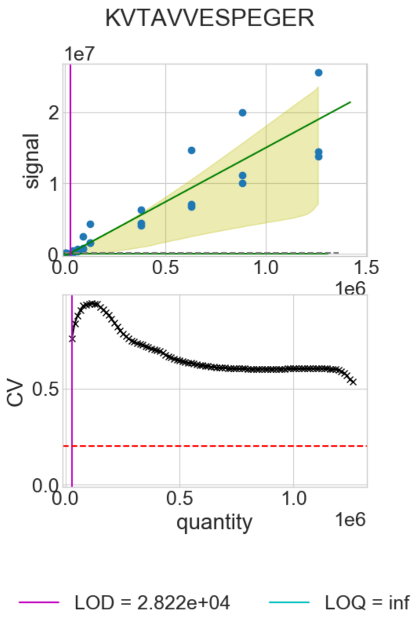


Figure 15: Calibration curve model outputs for Pma1 peptide KVTAVVESPEGER.

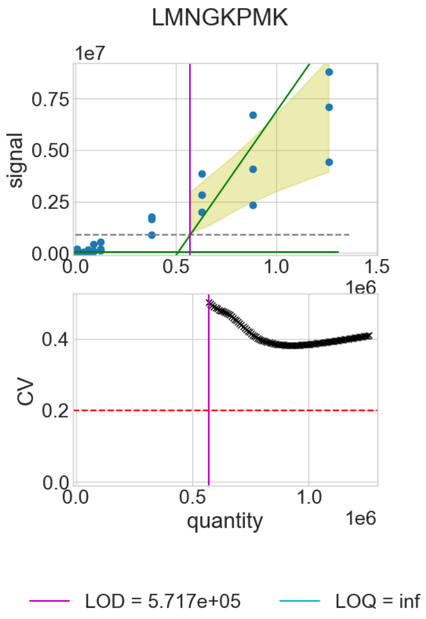


Figure 16: Calibration curve model outputs for Pma1 peptide LMNGKPMK.

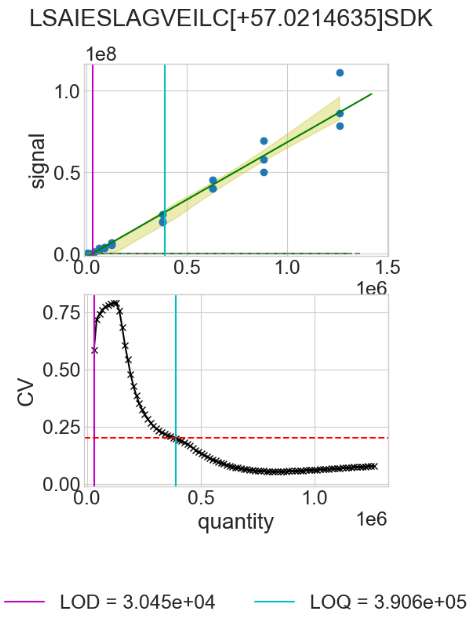


Figure 17: Calibration curve model outputs for Pma1 peptide LSAIESLAGVEILC[+57.0214635]SDK.

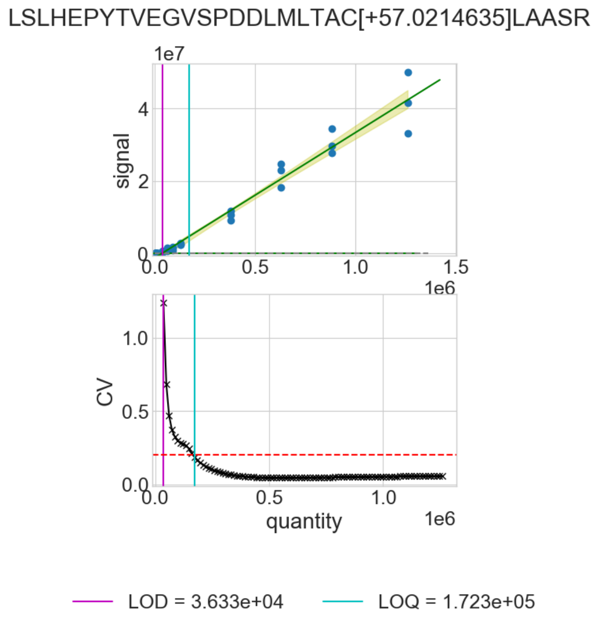


Figure 18: Calibration curve model outputs for Pma1 peptide LSLHEPYTVEGVSPDDLMLTAC[+57.0214635]LAASR.

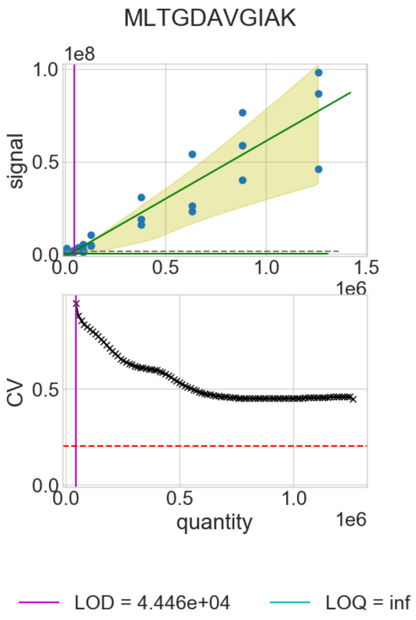


Figure 19: Calibration curve model outputs for Pma1 peptide MLTGDAVGIAK.

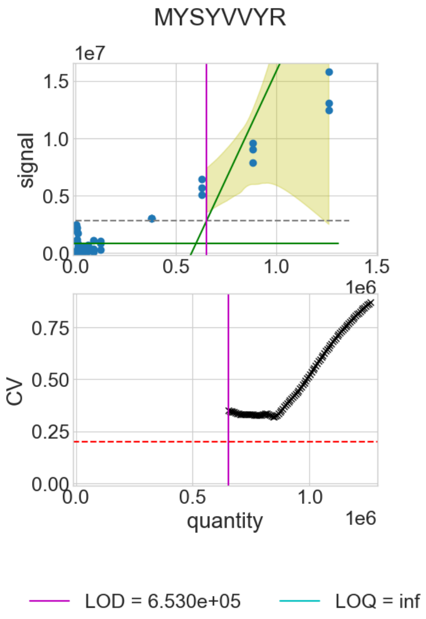


Figure 20: Calibration curve model outputs for Pma1 peptide MYSYVVYR.

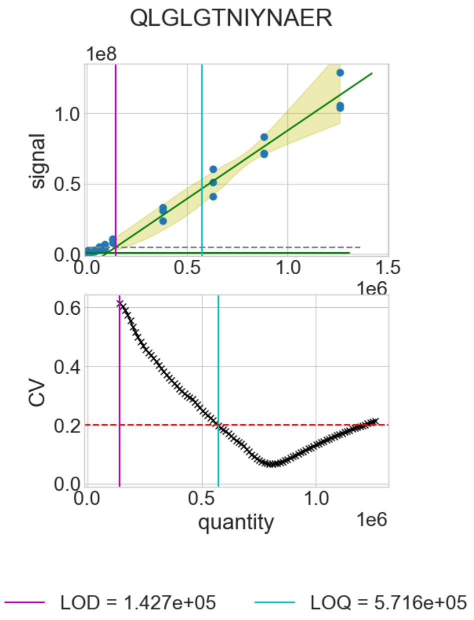


Figure 21: Calibration curve model outputs for Pma1 peptide QLGLGTNIYNAER.

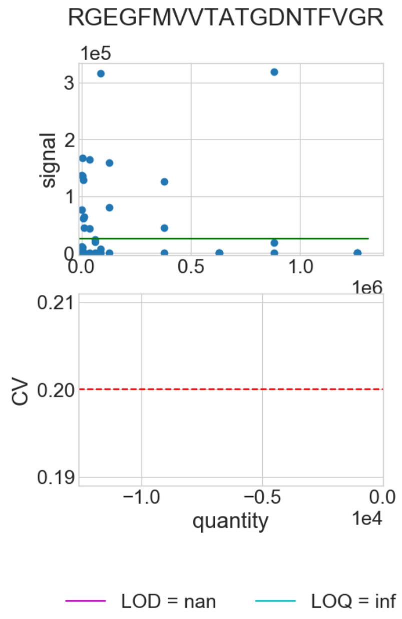


Figure 22: Calibration curve model outputs for Pma1 peptide RGEGFMVVTATGDNTFVGR.

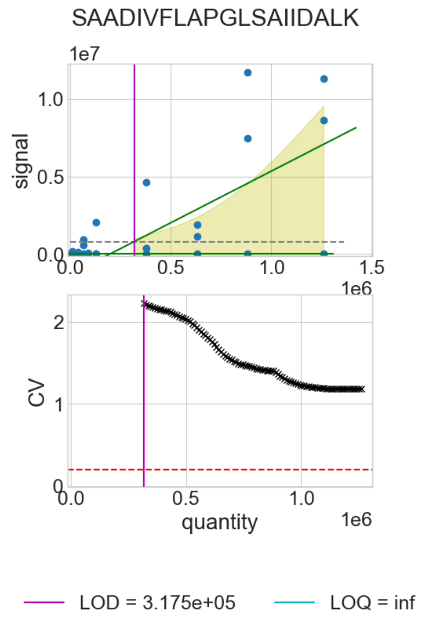


Figure 23: Calibration curve model outputs for Pma1 peptide SAADIVFLAPGLSAIIDALK.

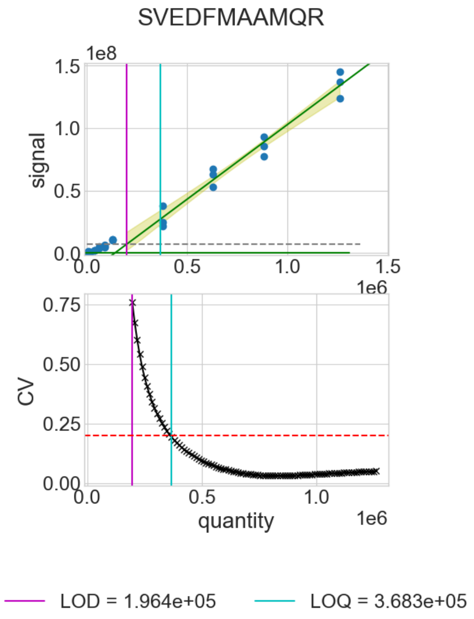


Figure 24: Calibration curve model outputs for Pma1 peptide SVEDFMAAMQR.

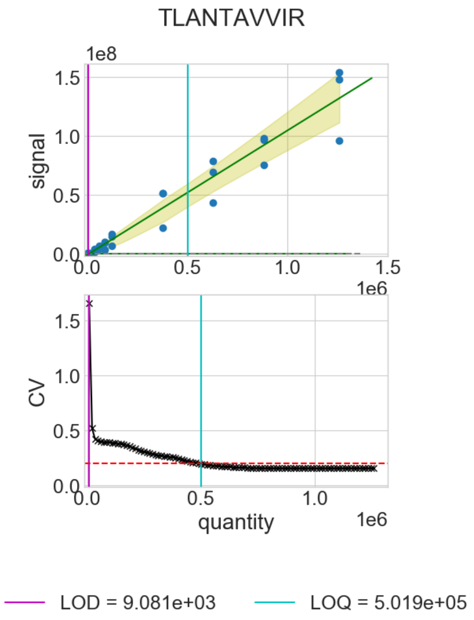


Figure 25: Calibration curve model outputs for Pma1 peptide TLANTAVVIR.

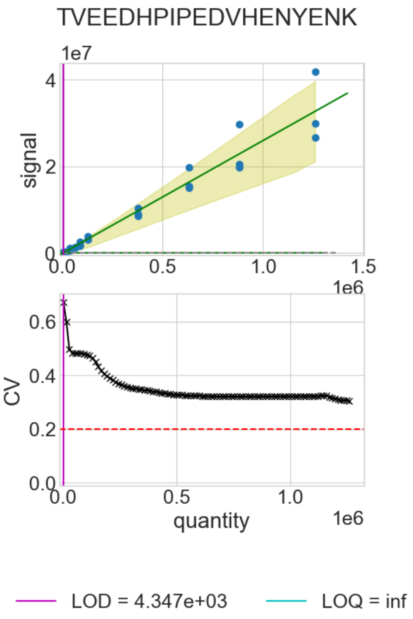


Figure 26: Calibration curve model outputs for Pma1 peptide TVEEDHPIPEDVHENYENK.

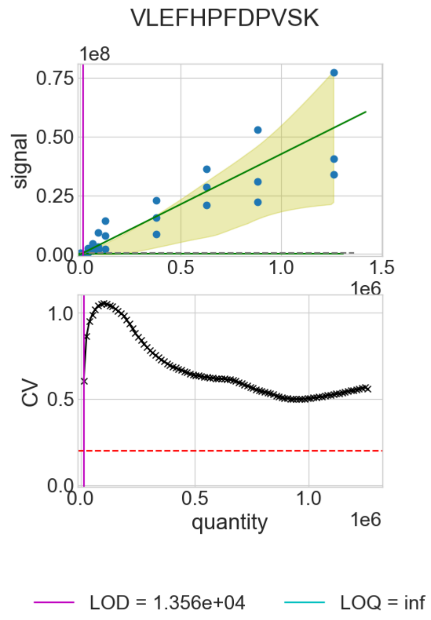


Figure 27: Calibration curve model outputs for Pma1 peptide VLEFHPFDPVSK.

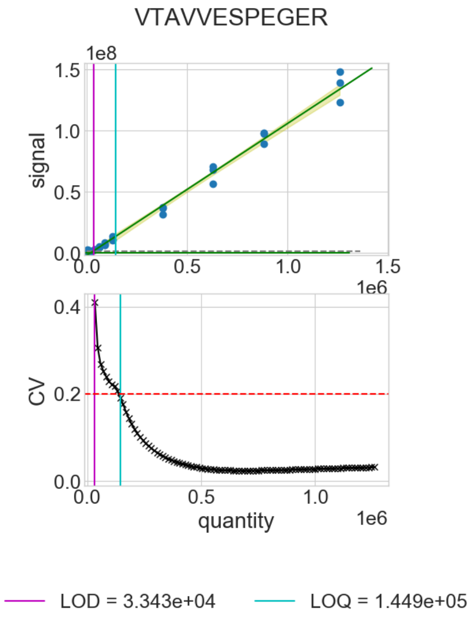


Figure 28: Calibration curve model outputs for Pma1 peptide VTAVVESPEGER.

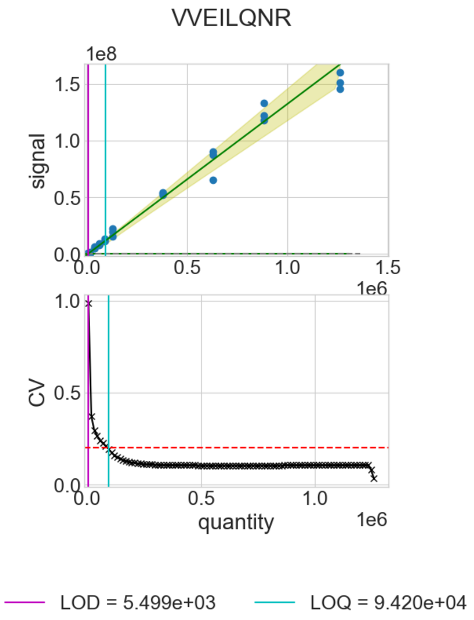


Figure 29: Calibration curve model outputs for Pma1 peptide VVEILQNR.

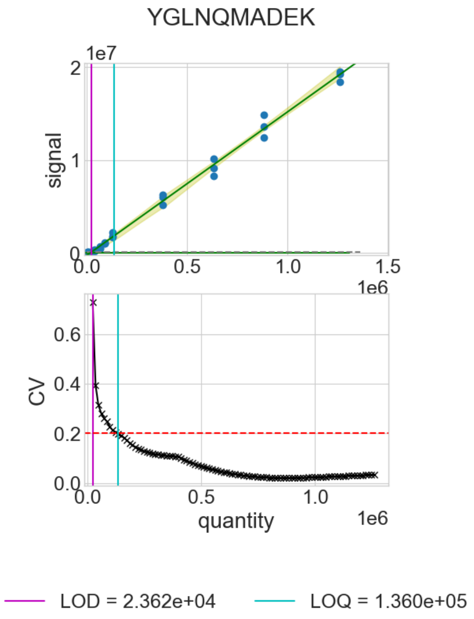


Figure 30: Calibration curve model outputs for Pma1 peptide YGLNQMADEK.

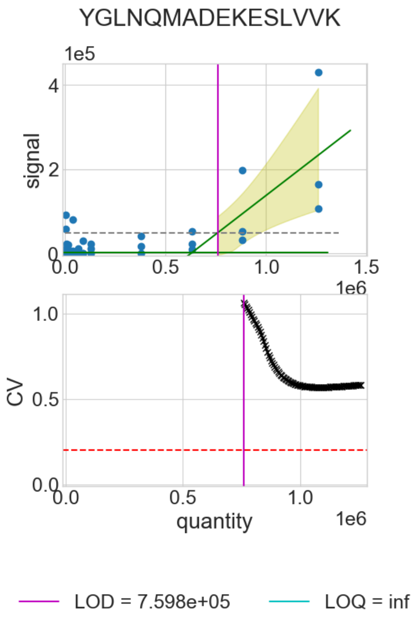


Figure 31: Calibration curve model outputs for Pma1 peptide YGLNQMADEKESLVVK.
